# Chemotherapy in Cases of *Trypanosoma cruzi* Reinfection: Assessing for Chagas Disease Endemic Areas

**DOI:** 10.1101/2021.08.27.457957

**Authors:** Mônica Cardozo Rebouças, Marcos Lázaro da Silva Guerreiro, Anderson Aires Eduardo, Sonia Gumes Andrade

## Abstract

People living in endemic areas of Chagas disease are submitted to multiple infections during their lives. This is an important factor in the development and morbidity of the disease. In the present investigation, evaluate the treatment outcome of triple infection mice with 21SF clones compared to the parental strain and with clones were investigated. Mice were infected and divided into groups: G1, infected with 21SF strain; G2, infected with 3 clones’s 21SF strain; and G3, infected with each clone alone. Subsequently, the groups were subdivided in treated and untreated controls. After the treatment the mice were euthanized. Serological tests and parasitological tests were performed. Sections of the heart and skeletal muscle were collected, fixed and then processed for the histopathological study in sections stained with Hematoxilin end Eosin. Parasitological tests for animals treated have shown positive results that varied from 25 to 66.7% and the serology titers varied from 1:10 a 1:280 in treated mice. Cure rates ranged from 11.1 to 30.8%. Histopathological examination revealed that treated animals presented clear reduction of lesions in myocardium and in skeletal muscle. Animals subjected to multiple infections have low rates of cure and worsening of tissue lesions.

## 1 INTRODUCTION

Chagas Disease (CD) is considered a public health and economy problem, which is considered one of the most important parasitic infections in Latin America (Pérez-Molina, J A; Molina, 2018). Moreover, the disease is considered not eradicable, since there no viable interruption of transmission throughout its wide range of hosts and reservoirs (Bern, 2015; Pérez-Molina and Molina, 2018). Besides, strains of *T. cruzi* represent complex multiclonal populations, which can be homogeneous or heterogeneous with predominance of some clonal lineage. It is recognized a wide biological variability among *T. cruz*i strains and several classification systems (Zingales et al., 2012).

As polyclonal strains, clones may present different degrees of virulence and differential responses to chemotherapeutic agents. For example, clones of the same strain 21SF, isolated from an endemic area of Chagas disease have a diversified behavior in relation to benznidazole (BZ) treatment (Andrade at al., 2006; Campos et al., 2005).

Infected individuals residing in endemic areas of CD are subject to successive reinfections by *T. cruzi*, provided that they remain in the same socioenvironmental conditions where vectors are present (Torrico et al., 2006). Such situations have been reported as critical for disease progression, affecting levels of cardiac lesions (Bustamante et al., 2007).

Considering that multiple infections can affect the dynamics of CD progression in a host organism, researches on this pathogenic context can improve the current understanding of chemotherapy under realistic endemic scenarios. In the present study we explore experimentally the response of organisms submitted to treatment with BZ after multiple infections with different clonal lineages from a same strain of *T. cruzi*. Specifically, we conducted experimental CD infections in mice using three clones obtained from the 21SF strain and evaluated the response to chemotherapy.

## 2 MATERIALS AND METHODS

### 2.1 Strain of *T. cruzi*

The 21SF strain of *T. cruzi*, isolated from acute human case of São Felipe municipality, state of Bahia, Brazil, was used in the present study. This strain is classified as biodeme type II (Andrade, 1974), zymodeme II (Miles et al., 1980) and *T. cruzi II* (Anonymous, 1999). The parental strain was maintained in laboratory by successive passage into mice and inoculated into the study group.

### 2.2 Cloning of the strain

Clonal lineages were established using micromanipulation, in which a single parasite were obtained from mice at 30th day of infection with the parental strain (Dvorak, 1985). Citrated blood collected was centrifuged at 900 g and the parasites in the plasma were counted in a Neubauer chamber after dilution in phosphate buffered saline (PBS), pH 7.2. A volume of 1 mL was distributed into multiwell microtitre culture plates and examined with an inverted microscope. Single trypomastigote forms were isolated and intraperitoneally inoculated into suckling mouse (8-days-old). From 10-30 days after the inoculation, the peripheral blood was examined for the presence of parasites, which were then defined as clones and classified as high, medium and low virulence (21SF-C8, C7 and C8, respectively). From positive animals, successive passages into suckling mice were performed for each clone. The clones were maintained in cryopreservation in liquid nitrogen at -196ºC and were thawed at 37ºC and immediately inoculated into Swiss mice (weighing 10-12g) to obtain the inoculum for the experimental groups.

### 2.3 Experimental groups

Swiss mice were used as model organism and were formed two experimental groups, Group I (triple infection), constituted of 100 mice infected successively with the three clones at intervals of 50 days (21SF-C6; 21SF-C7; 21SF-C8 - with 1 × 10^4^ trypomastigote blood forms for each clone); and Group II (single infection), constituted by 50 mice infected with 21SF parental strain and 25 mice infected with each clone separately, 21SF-C6, C7 and C8 (with 5 × 10^4^ trypomastigote blood forms for each ‘clone group’ infection).

### 2.4 Biological and infections parameters

Parasitemia was evaluated in five mice between 50th and 70th days after infection by microscopic examination of 5 µL of fresh tail blood, mounted in a glass slide and a cover-slip, were counted the parasites in 50 microscopic fields at 400X.

To perform the xenodiagnosis, five nymphs of III and IV stages of *Triatoma dimidiata* were used for each animal, conditioned in appropriate containers covered with nylon in the upper part. The animals were anesthetized and immobilized in sacks made with screens and placed in dorsal decubitus and the nymphs were then placed on the abdomen of each animal for a period of 30 minutes. After 30 and 60 days of xenodiagnostic preparation, the nymphs had their abdomen compressed to obtain the biological content (feces), which was diluted in PBS and examined in 50 microscopic fields, with a magnification of 400X, to detect parasitic forms.

The blood was collected from each mouse for haemoculture (0.5 mL), after euthanasia of the animal with negative parasitemia, were seeded in a test tube containing Warren medium (Warren, 1960) and maintained at 37ºC. For the determination of positivity, microscopic examinations of cultures with 30 and 45 days of culture were done.

The mortality was daily evaluated and recorded as the percentage of survivors during the experiments.

### 2.5 Treatment

Benznidazole (N-benzyl-2-nitro-1-imidazoleacetamide) was administered by intragastric gavage suspension at a dose of 100 mg/kg/day, with a total of 60 doses starting on the 14th postoperative day in the single infection groups and, in the triplicate infected group, thirty days after the third infection.

### 2.6 Histopathological study

Complete autopsies were performed, and several organs were fixed in 10% buffered formalin. Were collected sections of heart and skeletal muscle, fixed in formaldehyde/Millonig, included in paraffin and so obtained sections of 5 μm staining in hematoxylin and eosin (H & E) for histopathological study. The histopathological study was performed on mice infected with *T. cruzi*, comparing the intensity of fibrotic-inflammatory myocardial and skeletal muscle lesions between the non-treated groups of infection and the groups treated with BZ. A qualitative analysis was performed where the intensity of the inflammatory lesions was classified as discrete lesion, corresponding to scarce and diffuse mononuclear infiltrate and/or small foci of mononuclear infiltration; moderate lesion, represented by diffuse mononuclear infiltrate more pronounced than in degree and localized inflammatory foci, with focal myocyte lesion; intense lesion corresponding to dense diffuse infiltrate and/or extensive and confluent focal infiltrates and myocyte necrosis.

### 2.7 Indirect immunofluorescence serological test

This assay was performed according to Camargo (1966). The reaction was performed with different serum concentrations ranging from 1:10 to 1:1280 in multiwell slides using as antigens culture forms of *T. cruzi* and 10 µL of anti-mouse IgG fluorescein conjugated (Sigma) as specific antibodies. The slides were analyzed on a Zeiss epifluorescence microscope of with halogen lamp. Fluorescent reactions in sera were considered positive from ≥ 1:40 dilution, the cutoff point recommended by the Health Ministry (Dias et al., 2016).

### 2.8 The cure rates

It was established by combining the results of the parasitological tests and the immunofluorescence titers. Cured animals were non-positives in any of the cited tests.

The animals with a single and triple infection with each clone were sacrificed after 5 months of treatment. The surviving animals from the groups were euthanized by exsanguination under Ketamine and Xilasin anesthesia in PBS at a ratio of 1:1:1.

## 3 RESULTS

### 3.1 Parasitaemia and Cumulative mortality

The analysis of parasitemia and mortality rates showed that mice without treatment were more vulnerable to *T. cruzi* infection than mice treated with BZ. Parasitemia suppression was observed in animals submitted to chemotherapy (Figure 1; Table 1). Mice submitted to triple infection (Figure 1a) showed a parasitemic curve with peak in third week of first infection (with C6 clone) and a peak in first week after the second infection (with C7), i.e., the seventh week after first infection. We observed no peak after third infection (with C8) and after the treatment. The negative parasitemic was observed in tenth week of C8 infection (i.e., the week 25th after first infection). Mice under single infection showed no peak in parasitemic curve when submitted to chemotherapy, despite the clone used for infection (dotted lines in Figures 1c, 1e and 1g). In these cases, negative parasitemia was observed in the fourth week of infection for C6 and C8 and in the fifth week for C7. Non-treated mice display parasitemias with parasitemic peak between third- and fourth-week infection for C6, third week for C7 and fourth and sixth week for C8 (continuous lines in Figures 1c, 1e, and 1g). Negative parasitemia were observed at sixth week for C6, seventh week for C7 and C8. In the infection with 21SF strain, mices treated showed a parasitemic curve with no peak, whereas the non-treated control mice had a parasitemic peak in second week after infection. Negative parasitemia was observed in sixth week after infection for treatment group and tenth week for non-treated group (Figure 1i).

**Fig. 1:**
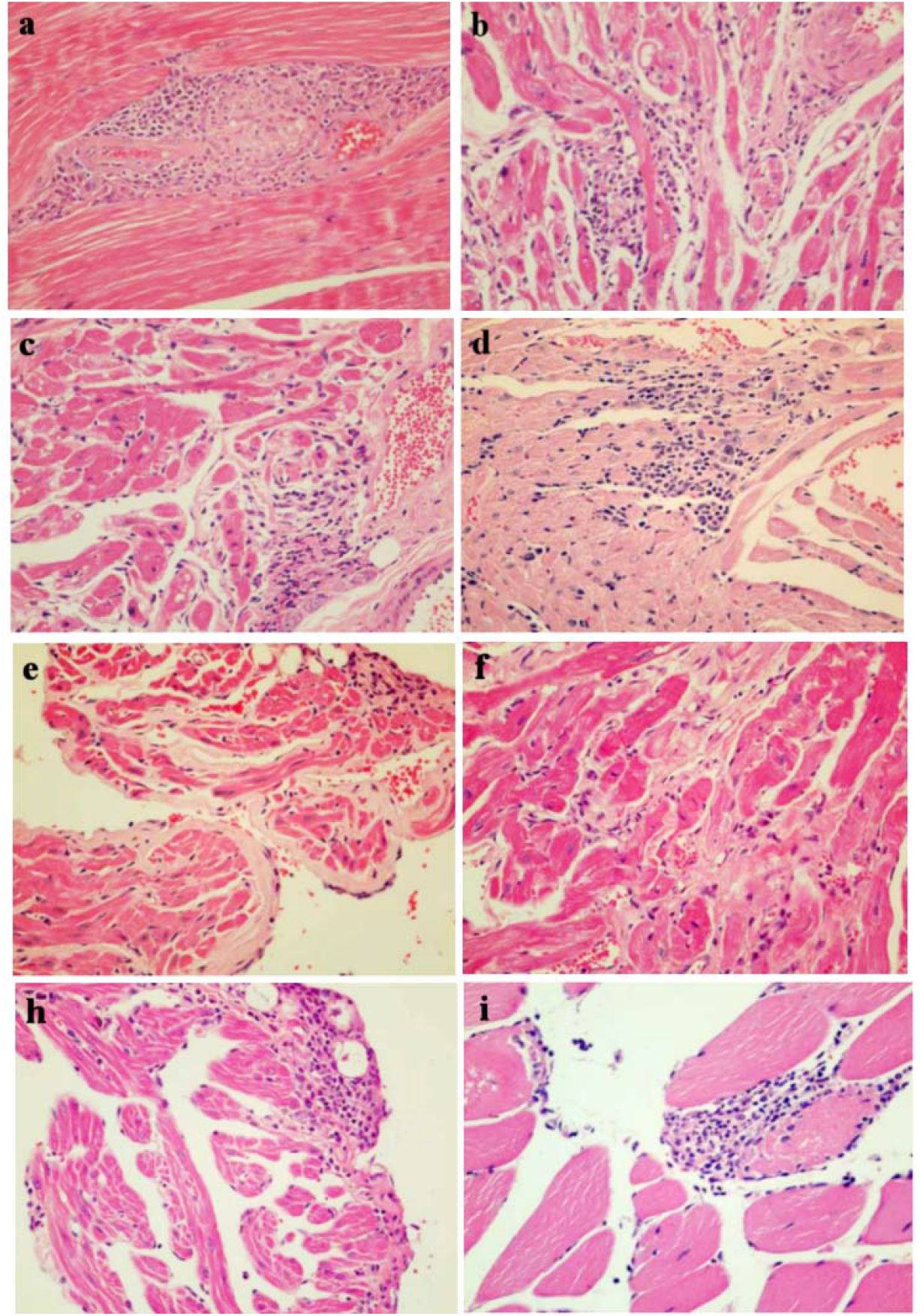
Heart and skeletal muscle sections of non-treated (a, c, e, g) and benznidazole (BZ) treated mice (b, d, f, h) stained with H&E (400X). Single infection with 21SF strain (a, b), C6 (c, d), C7 (e, f) and C8 (g, h). (a): skeletal muscle with focal myositis with perivascular inflammatory process. (b): myocardial section with focal and diffuse inflammatory lesions with areas of necrosis. (c): focal perivascular inflammatory infiltrate and focal subepicardial infiltrates in the myocardium with matrix changes. (d): atrium with discrete diffuse inflammatory infiltrate and area of moderate focal infiltrate. (e): section of the atrium with diffuse interstitial collagen deposits, predominating in the subepicardial region and focal inflammatory infiltrate. (f): myocardium with discrete diffuse interstitial infiltrate in the atrium with moderate collagen deposits. (g): myocardium with diffuse and discrete mononuclear inflammatory infiltrate in the atrium and moderate subepicardial focal infiltrate. (h): skeletal muscle presenting focal area of muscle fiber necrosis with neutrophil polymorphonuclear infiltrate.

**Fig 1.**
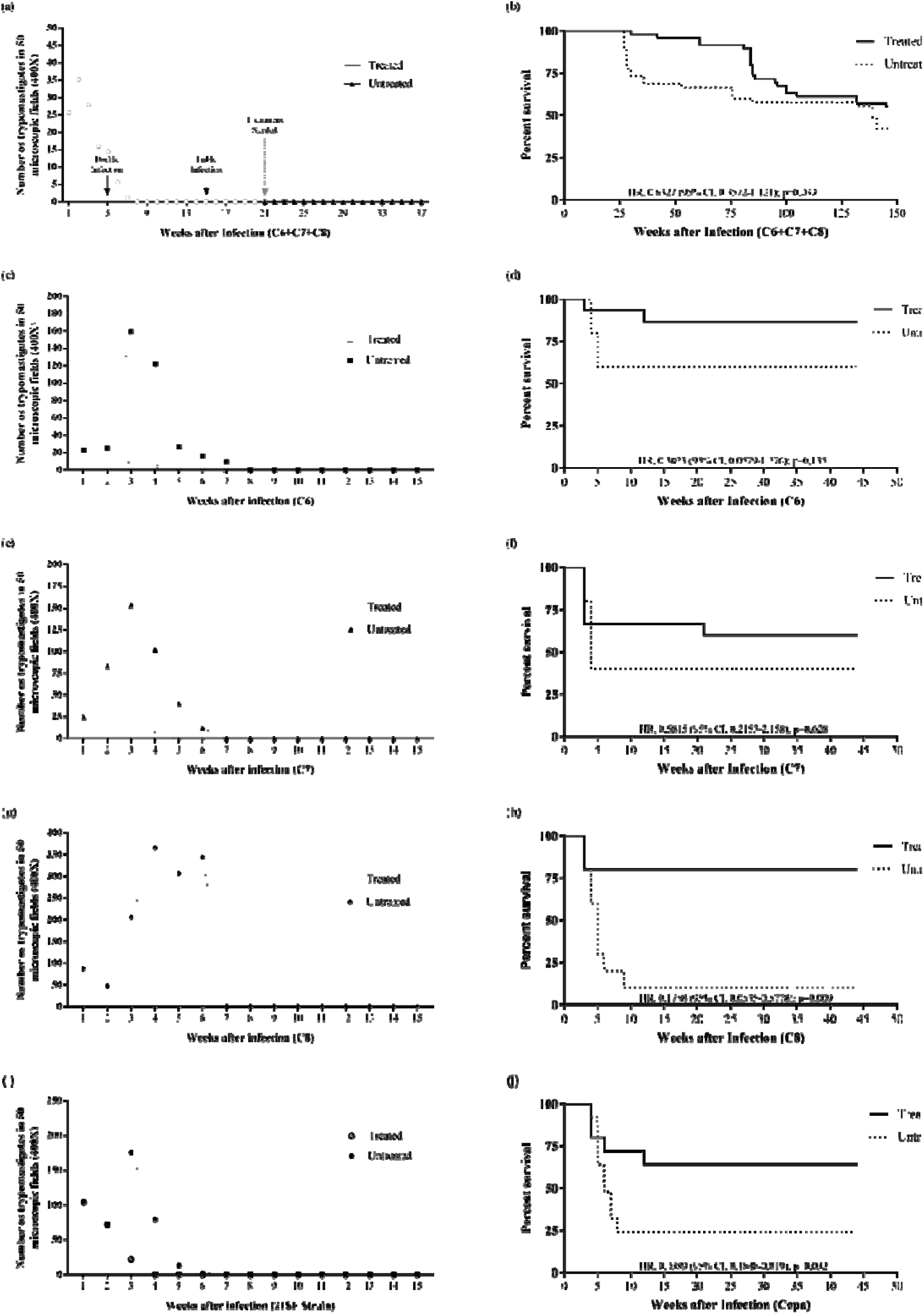
Median parasitemia curves in mice infected with clones of *T. cruzi* strain 21SF, showing the response of the group treated with benznidazole (BZ) (100 mg/kg/day) and of the control group. Parasitemia is shown in the graphics on the left, i.e., (a), (c), (e), (g), (i), being survival rate shown in the graphics on the right, i.e., (b), (d), (f), (h), (j). (a) and (b): triple infection with C6, C7 and C8 clones. (c) and (d): single infection with C6 clone. (e) and (f): single infection with C7 clone. (g) and (h): single infection with C8 clone. (i) and (j): single infection with 21SF strain. Survival curves were generated using the Kaplan-Meier method, Log-rank (Mantel Cox).

The mortality rates in the treated animals ranged from 13.3% for the C6-infected mice to 60% for the C7-infected group and 20% for the C8-infected mice. Mortality of the mice for the non-treated control groups was 40% for clone C6, 60% for clone C7 and 90% for clone C8 (Table 1).

**Table 1.** Analysis of parasitemia and mortality in Swiss mice infected with *Trypanosoma cruzi* and treated with benznidazole (BZ) or left non-treated.

| Experimental group |  | Highest<br>parasitemia level<br>(10<sup>4</sup>parasites/ml)<sup>a</sup> | *p* * | Week of<br>highest<br>parasitemia | Mortality<br>(%) | *p* ** |
| --- | --- | --- | --- | --- | --- | --- |
| **Triple**<br>**infection** | Treated | 0 | > 0,99 | - | 44.9 | 0,305 |
|  | Non-treated | 0 |  | - | 57.8 |  |
| **C6** | Treated | 8 ± 5 | < 0,01 | 3 | 13.3 | 0,175 |
|  | Non-treated | 151 ± 34 |  | 4 | 40 |  |
| **C7** | Treated | 45 ± 17 | < 0,01 | 3 | 40 | 0,428 |
|  | Non-treated | 197 ± 56 |  | 3 | 60 |  |
| **C8** | Treated | 11 ± 8 | < 0,01 | 3 | 20 | 0,001 |
|  | Non-treated | 441 ± 125 |  | 5 | 90 |  |
| **21FS**<br>**strain** | Treated | 26 ± 15 | < 0,01 | 3 | 36 | 0,009 |
|  | Non-treated | 201 ± 63 |  | 3 | 76 |  |

<sup>a</sup> Values are means (± standard deviations).
\* Mann-Whitney test
\*\* Fisher's Exact Test.

### 3.2 Parasitological Cure’s Test

Five months after the end of BZ treatment the mice were submitted to the curing test. The group with triple infection was positive in 29.6% for parasitological test, 30.7% for group infected with C6, 66.6% for mice infected with C7, 25% for group infected with C8 and 50% for group infected with 21SF strain.

### 3.3 Indirect Immunofluorescence Test

The indirect immunofluorescence test (IFI) revealed positive results for 95.2% of treated animals, infected triple with clones, 69.2% infected with C6, 88.9% infected with C7, 83.4% with C8 and 87.5% with 21SF strain. For the control experimental group, IFI revealed positive results of 100% for each infection group. The minimal titer considered as positive was 1:40. Titers varied from 1:10 to 1:280 for treatment groups and non-treated showed positive IFI with titers varying from 1:80 to 1:280.

### 3.4 Cure rates

The mice euthanized 5 months after the end of the treatment, were evaluated cumulatively to the curing tests to which they were submitted. Cure rates ranged from 11.1 to 30.8% based on the combination of parasitological and serological tests (Table 2).

**Table 2.**
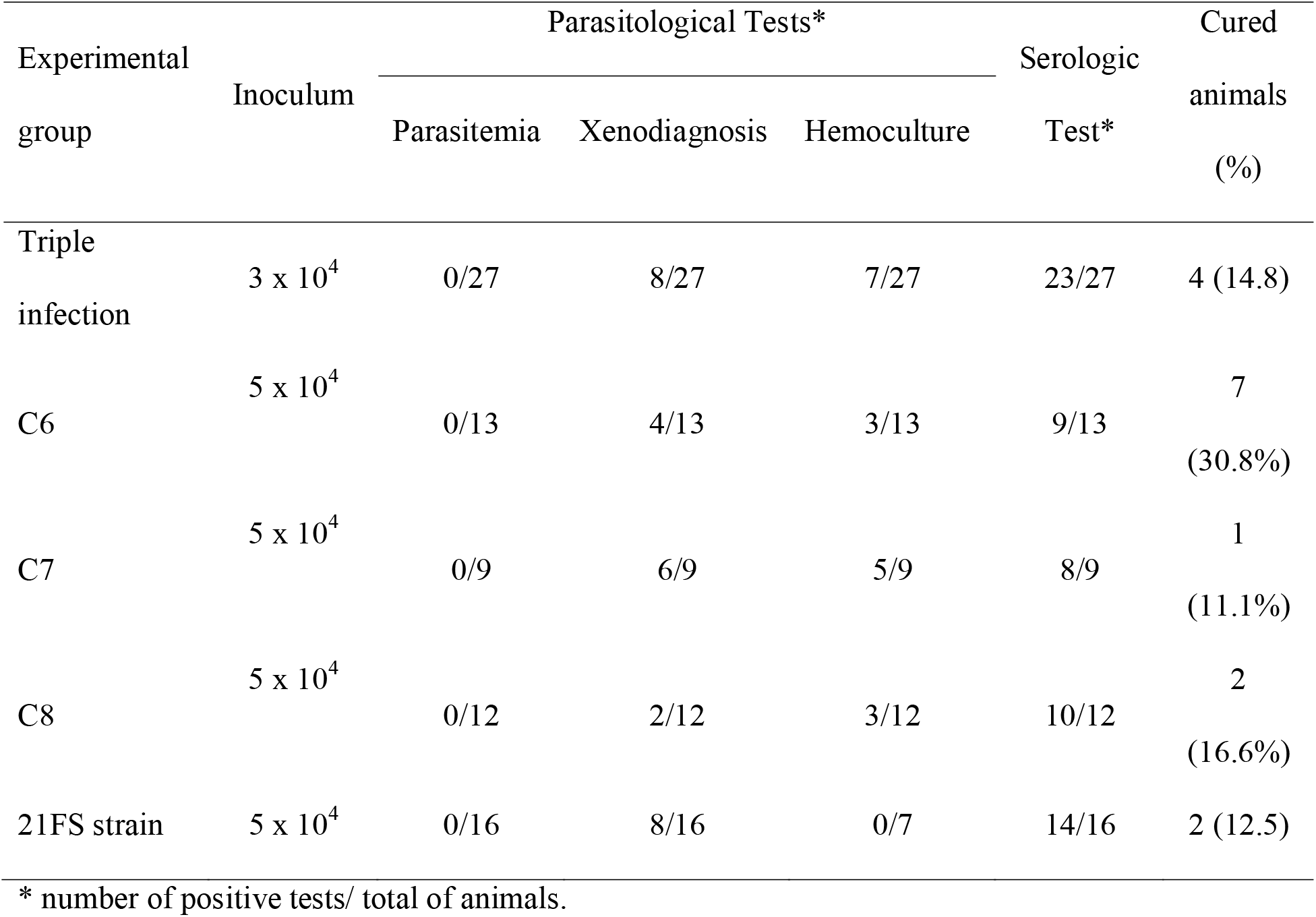
Cure rates of mice infected with *Trypanosoma cruzi* and treated with benznidazole (BZ).

### 3.5 Histopathology

In the histopathological study, we research the effect of treatment on inflammatory and necrotic lesions involving the myocardium and skeletal muscle in chronically infected mice (treated and non-treated). The infection with the parental strain determined in the chronic phase mild to moderate lesions, predominant in atria; the chronic lesions observed in non-treated and treated animals were similar (Figure 1a, 1b). The histopathological changes observed in the chronic phase of infection with the C6 clone were identical to those seen with the parental strain. The chronic lesions observed in non-treated and treated animals were similar (Figure 1c, 1d). In the C7 infection the lesions were identical to those observed with the parental strain. In both non-treated and treated animals, lesions of low intensity, limited to the atria, are observed; diffuse, moderate interstitial fibrosis occurred only in the non-treated. Decreased diffuse infiltrate and fibrosis were observed in treated animals (Figure 1e, 1f). In C8 infection, in the sole non-treated, acute-phase surviving mouse, there were mild to moderate inflammatory lesions in myocardium and skeletal muscle, like those observed in the group infected with the parental strain. In the mice treated with BZ, mild to moderate inflammatory lesions and mild interstitial fibrosis were observed, as well as focal alterations of muscle fibers with necrosis and hyalinization (Figure 1g, 1h).

In the triple infection, for controls, unlike in the other groups, we observed a compromise in both the atria and ventricles, with intense chronic myocarditis, and diffuse interstitial fibrosis. Skeletal muscle is much compromised, sometimes with very intense and diffuse lesions, with arthritis and intense perivascular interstitial lesions (Figure 2a-d). In the treated cases there was a clear reduction of the inflammatory lesions in both myocardium and skeletal muscle (Figure 2e-h).

**Fig. 2:**
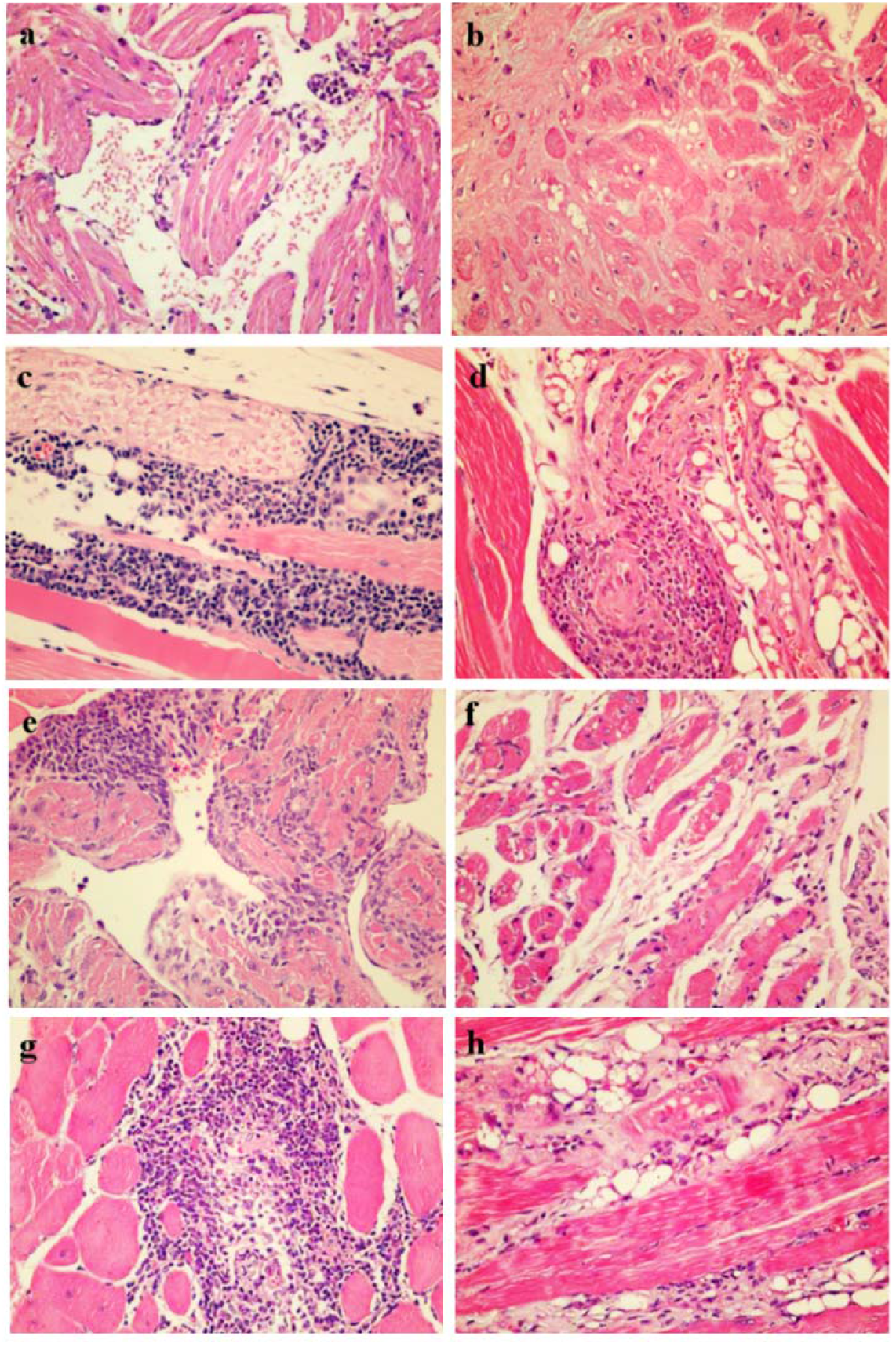
Heart and skeletal muscle sections of mice triple infected with C6, C7 and C8 clones, non-treated controls (a, b, c, d) and treated with benznidazole (BZ) (e, f, g, h) stained with H&E (400X). (a): section of the myocardium with diffuse and discrete mononuclear infiltration and subepicardial focal infiltrates. (b): ventricle section showing cardiac fibers with homogeneous cytoplasm and accentuated eusinophilia (hyaline necrosis) and presence of dense and diffuse interstitial matrix deposit. (c): section of the skeletal muscle presenting intense interstitial infiltration involving a vascular-nervous bundle and destruction of muscle fibers. (d): Vascular bundle with arthritis and dense periarteriolar infiltrate in skeletal muscle. (e): atrium with intense focal and diffuse mononuclear infiltrate. (f): myocardium with diffuse and moderate interstitial infiltrate and discrete matrix deposit. (g): area of focal destruction of muscle fibers replaced by intense mononuclear infiltrate (h): muscle fibers with vacuolization, replaced by interstitial inflammatory infiltrate and adipose infiltration.

## 4 DISCUSSION

In the present study, we evaluated the response to Benzonidazole (BZ) chemotherapy in mice with triple infection by *T. cruzi* 21SF strain, *T*.*cruzi II*. Considering the resistance to treatment in different strains, strain 21SF demonstrates an intraspecific variability in the response to chemotherapeutic agents (Andrade et al., 2006; Campos et al., 2005). Here, we observed low susceptibility of the 21SF strain to the chemotherapeutic, 12.5% cure, as well as the isolated clones (11.1 - 30.8%). This low susceptibility may be due to the genotypic and phenotypic clonal diversity of *T. cruzi*, probably due to the evolution of multiple lineages, as well as the addition of the serological test as a cure criterion (Andrade, 1999; Guerreiro et al., 2015; Romanha et al., 2010). In addition, 21SF strain was classified as moderately susceptible and individual evaluation may be required because of the wide range of biological responses found in mice (Andrade et al., 2006). The data presented here corroborate results found by other studies involving chemotherapy (Camandaroba et al., 2003; Campos et al., 2005) when evaluating the response of the strain and of five clones demonstrating that the susceptibility is related to multiple factors, such as parasite resistance, immunoregulatory aspects of number of serial passages.

The group with the triple infection were treated only in the chronic phase of the disease, with a cure rate of 14.8%. In the chronic phase of infection, the efficacy of BZ is very low (Pérez-Molina, J A; Molina, 2018) and other studies have shown natural resistance to chemotherapy of T. cruzi strains and clones (García-Huertas et al., 2017; Toledo et al., 2003). However, some researchers have suggested that the ineffectiveness is related to the pharmacokinetic properties of the drug, such as the relatively short half-life or limited tissue penetration, and the resistance to treatment of some strains (Boscardin et al., 2010; Lamas et al., 2006).

The results of the susceptibility of the clones of strain 21SF treated in the acute phase, showed a variation of 11.1 to 30.8% in the cure rates, through the parasitological and serological tests. Other clones isolated from this strain showed differences in their response to BZ (Campos et al., 2005), although they maintained the biological standard and the standard in restriction fragment length polymorphism (RFLP) analysis. However, demonstrating the efficacy of the treatment is complex because of inherent difficulties in establishing cure criteria and the limitations of the methods available to detect the cure of *T. cruzi* infection (Urbina, 2015).

Current treatment of Chagas disease is based on two drugs, Nifurtimox and Benznidazole (nitroheterocyclic compounds), developed empirically for more than 40 years (Urbina, 2018). These drugs are more effective in the acute phase of Chagas disease and in the chronic phase becomes low and with variable efficacy. In addition, side effects can lead to the interruption of the treatment in about 10-30% of the cases (Urbina, 2018, 2015). These data agree with our findings of treatment in the chronic phase. Moreover, previous studies also have been based on parasitological tests, including parasitemia, xenodiagnosis, blood culture and subinoculation in newborn mice, combined with specific serological titers and establishing a limit of 1: 20 (Andrade et al., 1985). These joint assays are currently used as parasitological cure criteria in humans and experimental animals.

In the data presented here, the presence of anti-*T. cruzi* in BZ treated animals was 82% for all single infection groups and 85% in the triple infection group. In the untreated controls, positivity was 100% in all groups. The high serological titers detected in animals with single or triple infection after prolonged treatment with BZ can be explained by the presentation of antigens by dendritic cells present in the spleen (Portella and Andrade, 2009).

Pinto et al. (2013) in a clinical cohort study evaluating the response to Benzonidazole treatment in an acute phase patient in the State of Amazonia demonstrated that after treatment there is a significant decrease in circulating levels of IgG and IgM. These findings confirm the findings found when the immune response was investigated in six different strains of mice infected by different strains of *T. cruzi* (Andrade et al., 1985). Based on our findings and the related literature, we may suggest that our results are linked to complex immunoregulatory processes, most likely related to the involvement of cytokines, which may influence the expression of antibody isotypes during treatment infection and disease. In this way, we argue that the definition of conventional serological cure is often hampered because the serological tests can remain positive for a long time after the cure.

The results of our parasitological tests corroborate the hypothesis of the average susceptibility presented by this lineage, with cure rates varying from 33.3 to 84%. But when taken together with serological titers, cure rates range from 11.1% to 30.8%. Caldas et al. (2008) demonstrated that the induction of resistance of *T. cruzi* strains to BZ may also occur during long-term infection in the vertebrate host and undergo successive passages in mice in the absence of treatment. This may explain the absence of cure observed in individuals with chronic Chagas’ disease treated with BZ.

The diagnosis of chronic infection and the cure test are made by serological tests, more commonly with the use of an immunoenzymatic assay (ELISA) or immunofluorescence assay (IFA), which must present a persistent and progressive reduction or reduction of the three titration dilutions (Bern, 2015). Polymerase Chain Reaction (PCR) can be used as an auxiliary method for such analysis and its positivation represents therapeutic failure – not used in the limitation of the techniques (Dias et al., 2016).

Although Type II biodema strains, such as the 21SF strain, have high variability in their response to chemotherapy, their behavior may be considered dependent on the predominance of a resistant clone representative of the strains of the *T. cruzi II* taxa (Andrade, 1999). The resistance/natural susceptibility of T. cruzi to BZ has been described as an important factor to explain the low cure rates detected in chagasic patients (Chevillard et al., 2018; García-Huertas et al., 2017).

The resistance to chemotherapy presented by some clones studied may be associated with successive passages in guinea pigs and triatomines as a way of maintaining these isolates. Caldas et al. (2008) demonstrated that the induction of resistance of *T. cruzi* to BZ strains may also occur during long-term infection in the vertebrate host and undergo successive passages in mice in the absence of treatment. This may explain the absence of cure observed in individuals with chronic Chagas’ disease treated with BZ.

The group with triple infection had successive inoculum, with intervals ranging from approximately seven weeks, totalizing 3×10^4^ trypomastigote blood forms, before the beginning of the treatment. The groups corresponding to the single infection had an inoculum of 5×10^4^ trypomastigotes blood forms. This infection scheme had the objective of avoiding in the triple infection a total mortality of the mice submitted to infection.

The chronic phase histopathology, observed in the present study, in treated and untreated BZ controls, submitted to a single infection (control groups) with clones of 21SF strain, revealed lesions varying from mild to moderate, such as those found in the strain infection parental. The evaluation in the group submitted to triple infection showed exacerbation of the lesions, both in the myocardium and skeletal muscle. It has been demonstrated that successive inoculations with different strains of *T. cruzi* and clones can influence the mechanisms of immune response in the mouse, potentializing and intensifying the inflammatory response, leading to an increase in the lesions found in three-infected mice (Andrade, at al., 2006; Guerreiro et al., 2015). Studies the histopathology with two T. cruzi strains, the importance of the inoculum in the production of lesions was verified: the low inoculum can determine significant lesions, which vary from mild to moderate, besides being able to influence parasitemia, inflammation and cytokine production (Vazquez et al., 2015).

Previous data suggest that, from the serological and parasitological point of view, the cure rates determined by the treatment are highly variable in the chronic phase of the disease (Rassi et al., 2017). In addition, Bustamonte et al. (2007) demonstrated that reinfections in the acute phase of *T. cruzi* infection produced a more severe chronic phase and the prognosis of the disease.

## 5. CONCLUSION

The present results suggest that from the joint analyzes that the results in animals infected with single and triple infection, considering the parasitological and serological tests, have low cure rates. Infected triplicate animals presented worsening of myocardial and skeletal muscle inflammatory lesions when compared to the mice of the control group with single infection. These results suggest that patients living in endemic areas are subject to worsening of the pathological condition of the disease.

---

**TABLE 1.** Analysis of parasitemia and mortality in Swiss mice infected with *Trypanosoma cruzi* and treated with benznidazole (BZ) or left non-treated (± standard deviations).

| Experimental group |  | Highest parasitemia level<br>(10<sup>4</sup> parasites/ml)<sup>a</sup> | Week of highest parasitemia | Number of surviving/ total number of animal | Mortality (%) |
| --- | --- | --- | --- | --- | --- |
| Triple infection | Treated | 0 | - | 27/50 | 44.9 |
|  | Non-treated | 0 | - | 19/45 | 57.8 |
| C6 | Treated | 8 ± 5 | 3 | 13/15 | 13.3 |
| | Non-treated | 151 ± 34 | 4 | 6/10 | 40 |
| C7 | Treated | 45 ± 17 | 3 | 9/15 | 40 |
| | Non-treated | 197 ± 56 | 3 | 4/10 | 60 |
| C8 | Treated | 11 ± 8 | 3 | 12/15 | 20 |
| | Non-treated | 441 ± 125 | 5 | 1/10 | 90 |
| 21FS strain | Treated | 26 ± 15 | 3 | 16/25 | 36 |
| | Non-treated | 201 ± 63 | 3 | 6/25 | 76 |

<sup>a</sup> Values are means.

**TABLE 2.**
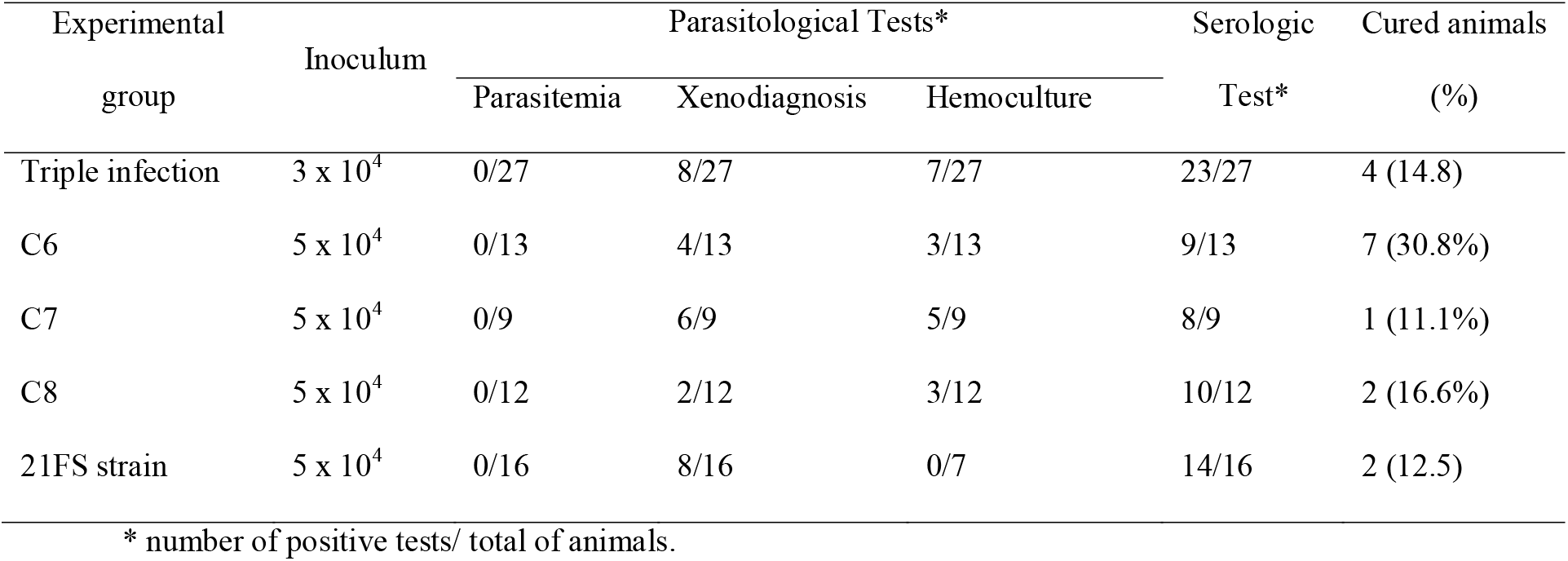
Cure rates of mice infected with *Trypanosoma cruzi* and treated with Benznidazole (BZ).

